# Biomolecular condensates are characterized by interphase electric potentials

**DOI:** 10.1101/2024.07.02.601783

**Authors:** Ammon E. Posey, Anne Bremer, Nadia A. Erkamp, Avnika Pant, Tuomas P.J. Knowles, Yifan Dai, Tanja Mittag, Rohit V. Pappu

## Abstract

Biomolecular condensates form via processes that combine phase separation and reversible associations of multivalent macromolecules. Condensates can be two- or multi-phase systems defined by coexisting dense and dilute phases. Here, we show that solution ions can partition asymmetrically across coexisting phases defined by condensates formed by intrinsically disordered proteins or homopolymeric RNA molecules. Our findings were enabled by direct measurements of the activities of cations and anions within coexisting phases of protein and RNA condensates. Asymmetries in ion partitioning between coexisting phases vary with protein sequence, condensate type, salt concentration, and ion type. The Donnan equilibrium set up by asymmetrical partitioning of solution ions generates interphase electric potentials known as Donnan and Nernst potentials. Our measurements show that the interphase potentials of condensates are of the same order of magnitude as membrane potentials of membrane-bound organelles. Interphase potentials quantify the degree to which microenvironments of coexisting phases are different from one another. Importantly, and based on condensate-specific interphase electric potentials, which are membrane-like potentials of membraneless bodies, we reason that condensates are mesoscale capacitors that store charge. Interphase potentials lead to electric double layers at condensate interfaces. This helps explain recent observations of condensate interfaces being electrochemically active.

## INTRODUCTION

One of the defining hallmarks of cells is the resting membrane potential created by differences in the permeabilities of ions ^1^. Thermodynamically, passive membrane potentials, also known as the reversal potential, are defined by the Gibbs-Donnan equilibrium ^2^. This equilibrium potential is set up by asymmetrical distributions of ions across semi-permeable membranes and is defined by the composition of lipid molecules and the proteins that are embedded in the membrane ^3^. The passive membrane potential is modulated by membrane-spanning ion channels, transporters, pumps, and exchangers ^4^. Systems comprising two or more coexisting phases but lacking membranes are also characterized by Donnan potentials because this potential is also set up by the Gibbs-Donnan equilibrium between phases ^2, 5^. Therefore, a membrane-like potential should also be realizable in inhomogeneous two- and multiphase systems that lack a membrane. This is relevant for biomolecular condensates ^6^, which are membraneless entities featuring coexisting phases ^7^.

Condensates form via composite processes that combine phase separation, reversible associations, and continuous phase transitions such as percolation ^8-11^or charge-driven complex coacervation ^12^. The latter process ^13^ comes in different flavors ^14^ and it can give rise to physiologically relevant ^10^ or *de novo* nuclear condensates ^15^. Phase separation is the underlying process that gives rise to two or more coexisting phases. Phase equilibria are established by equalization of species-specific chemical potentials and equalization of the osmotic pressure across the coexisting phases ^8^. Differences in macromolecular concentrations between dilute and dense phases are examples of interphase properties that arise from equalizing the chemical potentials of macromolecules ^10^. At equilibrium, the chemical potentials of each species in the solvent must also be equalized across coexisting phases. In physiologically relevant aqueous buffers, the most prominent non-aqueous components of solvent are monovalent cations and anions ^10, 16^. If equalization of chemical potentials gives rise to differences in activities of cations and anions across the coexisting phases (**Figure 1A**), then condensates will be characterized by interphase electric potentials (**Figure 1B, 1C**). Note that the term condensate refers to the two-phase system, and not just to the dense phase.

**Figure 1:**
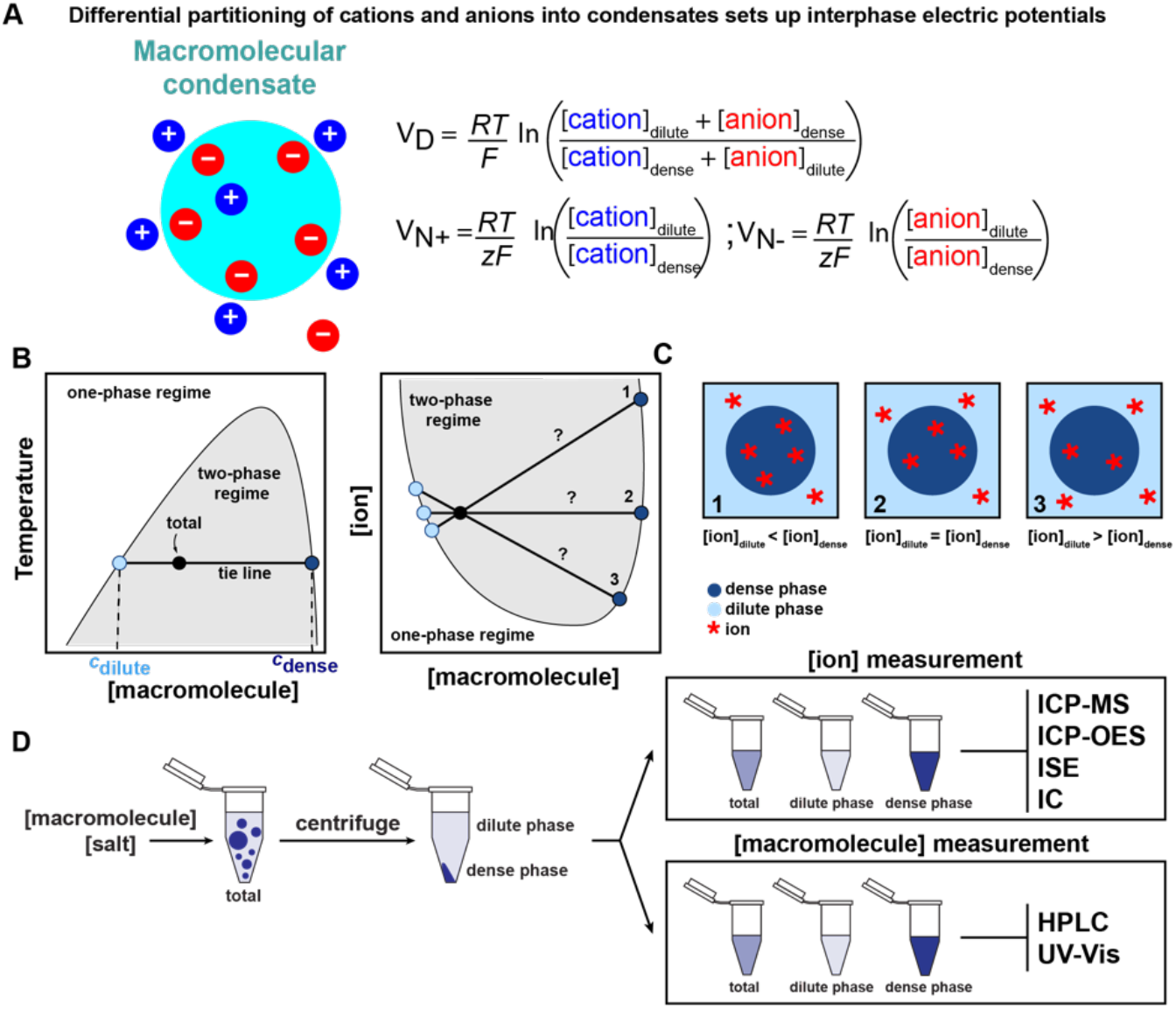
Differential partitioning of solution components will set up interphase electric potentials. **(A)** Phase separation gives rise to a macromolecule-rich phase (cyan) that coexists with a macromolecule-deficient phase (white background). If phase equilibrium is established by differential partitioning of cations and anions, then an interphase Donnan potential (V_D_) will be established. Likewise, differential partitioning of cations and anions will set up two sets of Nernst potentials, denoted here as V_N+_ and V_N-_, respectively. Direct measurements of the activities of cations and anions in coexisting dilute and dense phases can be used to quantify the magnitude and sign of V_D_ using the Goldman-Hodgkin-Katz equation, as well as V_N+_ and V_N-_ using the Nernst equation. Here, *R* = 8.314 J/K-mol is the ideal gas constant, *T* is the temperature of the system, and *F* = 9.645 × 10^4^ C / mol is the Faraday constant. **(B)** Example of a phase boundary of a system that undergoes temperature-dependent phase separation with upper critical solution temperature. If the conditions (temperature and concentration) for a given sample (black circle) fall within the two-phase regime (grey region), then the sample will separate into dense (concentration, *c*_dense_) and dilute phases (concentration, *c*_dilute_), respectively). The tie line connects the concentrations of components such as the macromolecule whose dense and dilute phase concentrations are constrained by equalization of chemical potentials. Here, a horizontal tie line indicates that the temperature is the same in the dense and dilute phases. **(C)** Tie lines can have slopes as can be seen for salt-dependent phase separation. Unlike horizontal tie lines, which indicate equal distribution of ions between phases (scenario 2), sloped tie lines indicate that the ion in question is preferentially included (scenario 1) or is preferentially excluded (scenario 3) from the dense phase. **(D)** Workflow of sample preparation for measuring protein and ion concentrations in the bulk solution, as well as dilute and dense phase samples. Concentrations of macromolecules were measured with HPLC and ultraviolet-visual (UV-Vis) spectroscopy. We directly measure the salt concentrations in the total sample as well as the coexisting dilute and dense phases to construct tie lines. To measure ion concentrations, we used inductively coupled plasma mass spectrometry (ICP-MS), ion selective electrodes (ISE), inductively coupled plasma atomic emission spectroscopy (ICP-OES) and ion chromatography (IC).

The Galvani potential is an interphase electric potential that quantifies the difference in electric potentials of the components between two points in the solution, one located in the dilute phase, and the other in the dense phase ^17^. In a two-phase system, the interphase Donnan potential, also referred to as the membrane potential of a membrane-bound system, is the Galvani potential that originates in differential partitioning of ionic species between the dilute and dense phases ^2, 5^. If activities of cations and anions in coexisting dense and dilute phases can be measured, then the Donnan potential, denoted as V_D_, can be quantified using the Goldman-Hodgkin-Katz equation ^18^ (**Figure 1A**). The Donnan potential quantifies the amount of charge that can be stored in or depleted from the two-phase system, depending on whether this potential is positive or negative.

The interphase Nernst potential is the Galvani potential that arises from asymmetrical partitioning of specific ionic species at equilibrium ^19^. Assuming an aqueous solution comprising one type of major cation and one type of major anion, one can estimate Nernst potentials denoted as V_N+_ and V_N-_ (**Figure 1A**) providing we know the concentrations of cations and anions in each of the coexisting phases ^20-22^. A Nernst potential ^21^ is a measure of the voltage needed to drive the flow of specific ions into or out of the condensate. The sign of the potential points to the directionality of cation versus anion flow. Hence, a Nernst potential can be interpreted as the voltage that must be applied across the two-phase system to balance any difference in activities of specific ionic species across the dilute and dense phases ^20, 21^.

We report results from direct measurements of the activities of monovalent cations and anions in coexisting dilute and dense phases of condensates formed by the intrinsically disordered prion-like low complexity domain (PLCD) derived from hnRNPA1 and variants thereof ^23, 24^, and nucleic-acid condensates formed by poly-rA RNA molecules ^25^. The measured activities were used to quantify interphase electric potentials of model biomolecular condensates in aqueous solutions comprising monovalent ions. We used a combination of high-performance liquid chromatography (HPLC) and ultraviolet-visual (UV-Vis) absorption spectroscopy for direct measurements of macromolecular activities in coexisting dilute and dense phases ^26^. Activities of cations in coexisting phases were measured using elemental analysis methods, specifically inductively coupled mass spectrometry (ICP-MS) ^27^ and inductively coupled optical emission spectroscopy (ICP-OES) ^28^. The ICP-based methods are highly sensitive, but they can only be deployed to measure cation activities. Accordingly, activities of anions in coexisting phases were measured using ion selective electrodes (ISE) ^29^ and ion chromatography (IC) ^30^ (**Figure 1D**).

Our measurements show the presence of condensate-specific interphase Donnan and Nernst potentials that are of the same order of magnitude as membrane potentials ^20-22^. The interphase electric potentials depend on macromolecular charge, the mean ion activity in a specific monovalent salt, and the identity of the ions that make up the monovalent salt solution.

## RESULTS

### Conceptual foundations underlying the measurements of interphase potentials

Consider two coexisting phases designated as A and B, respectively. Phase equilibrium is established by the equalizing chemical potentials for each of the species in solution. We make this point using an arbitrary component designated as *X*. The chemical potentials in the A and B phases are written as: 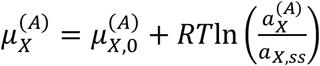 and 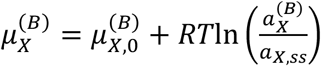, respectively. Here, 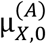 and 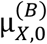 denote the standard state chemical potentials of species *X* in the two phases A and B. Note that 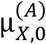 and 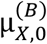 are partial molar free energy changes associated with transferring species *X* from a shared standard state into phases A and B. Therefore, they quantify the affinities of species *X* for phases A and B, respectively. If *X* is an ionizable species, then the standard state electrochemical potentials are rewritten as: 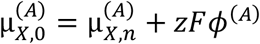 and 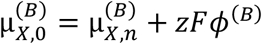. Here, the μ_*X,n*_ are standard state chemical potentials of the uncharged form of *X*. The contribution from the electrical potential is written in terms of *z*, which is the charge on *X, F* the Faraday constant, and *ϕ* the electric potential within the specific phase. The quantities 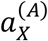 and 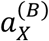 are activities of *X* in phases A and B. Note that the activity is the concentration of free *X* in each of the phases. If the activity coefficient is unity, then the activity and the concentration are identical. Note that the activities are referenced to *a*_*X,SS*_, which is the activity of *X* in the state that is used to estimate the standard state chemical potentials. We measure the activity *X* in each phase. Finally, *RT* is a measure of thermal energy. We set *R* = 1.987×10^−3^ kcal/mol-K and *T* = 298 K in our measurements.

Equalizing the chemical potentials of *X* across the coexisting phases results in: 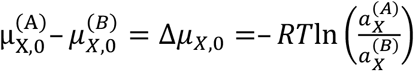. If the difference in standard state chemical potentials of species *X* is zero, then 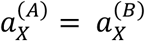. Conversely, if the difference in standard state chemical potentials of species *X* is non-zero, then at equilibrium we expect there to be a passive interphase gradient of species *X* because either 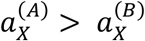 or 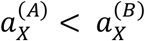. In our measurements, the species of interest *X*_1_, *X*_2_, …, *X*_n_ will be the macromolecule that drives condensation as well as the anions and cations that make up the solution. An imbalance of anions and cations will set up interphase electric potentials, and we quantify these potentials, using direct measurements as a function of salt concentration and salt type.

### Phase separation of A1-LCD gives rise to interphase electric potentials

We measured dilute and dense phase concentrations of proteins, cations, and anions in condensates formed by A1-LCD and three different variants of this protein (**Figure 2A, 2B**). Here, A1-LCD refers to the PLCD of the RNA-binding protein hnRNPA1 (**Figure 2A**). It has a net charge of +8 and its amino acid sequence features uniformly distributed aromatic residues interspersed by Gly, Ser, and ionizable residues (**Table S1**) ^23, 24, 31^.

**Figure 2:**
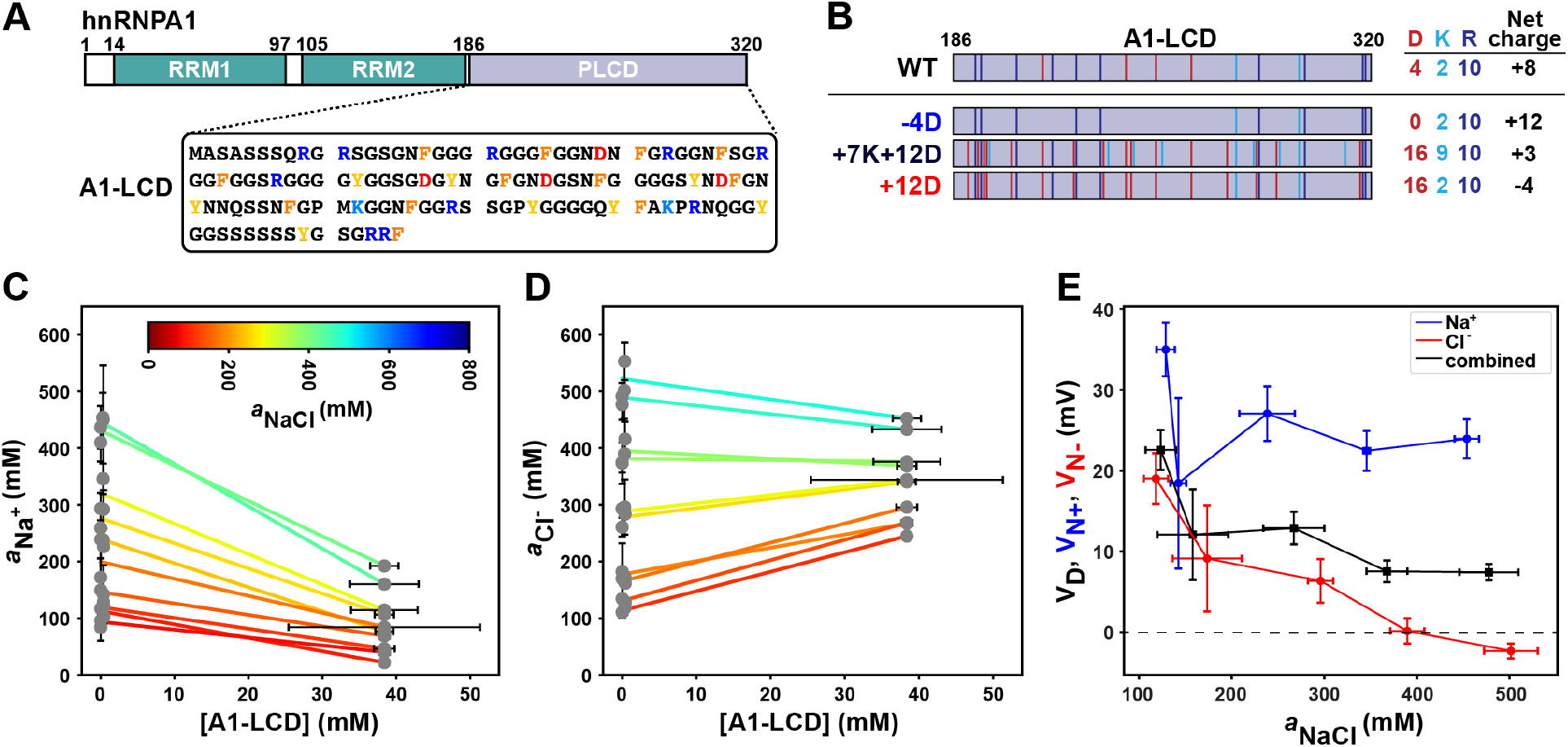
Na^+^ and Cl^-^ ions partition differentially across dense and dilute phases formed by A1-LCD. **(A)** Schematic of the sequence architecture of the RNA binding protein hnRNPA1. The sequence encompasses two RNA recognition motifs (RRM1 and RRM2) and an intrinsically disordered prion-like low complexity domain (PLCD) that we designate as A1-LCD. **(B)** We studied the phase behaviors of the wildtype (WT) A1-LCD and three other designed variants. These are defined by differences in net charge, achieved by replacing Asp residues with Gly / Ser (−4D) or replacing Gly / Ser residues with either Lys or Asp (+7K+12D or +12D). The figure shows quantification of the number of Asp (D), Lys (K), and Arg (R) residues, as well as the total net charge of each variant. Activities of Na^+^ **(C)** and Cl^-^ **(D)** were measured by ICP-MS and ISE, respectively. Measurements were performed for different total ion activities (see color bar in panel C). Tie lines were determined as linear fits of measured concentrations for dilute phase, the total sample, and dense phase. Note that sloped tie lines indicate differential partitioning of cations and anions between dense and dilute phases. The slopes of tie lines are negative for Na^+^ whereas the slopes for Cl^-^ become less positive as the total ion activity increases. The change in slope for Cl^-^ is thermodynamically linked to the extent to which Na^+^ is excluded. Tie lines are colored according to the measured total ion activities, as indicated by the color bar. Vertical error bars indicate the standard deviation across three technical replicates, and data are shown for two independent protein preparations (biological replicates). Dense phase concentrations are reported as the mean of all (ten) dense phase samples, and the horizontal dense phase error bars represent the absolute deviation from the mean. All measured salt concentrations were converted to activities. **(E)** Interphase Donnan and Nernst potentials are plotted against total ion activity. The dashed horizontal line is drawn to guide the eye and it intersects the ordinate at a value of zero. See **Figure S1** for additional data. Tie lines should not cross one another. However, some level of tie line crossing is evident in panels (C) and (D). This is a direct consequence of depicting a line in 3-space as a line on a plane. Indeed, as shown in supplementary movies, the lines in 3-space do not cross one another. See **Movie S1**.

A1-LCD forms coexisting dilute and dense phases in solutions with different concentrations of NaCl and with different types of monovalent salts (**Figure S1A**). We first measured the activities of A1-LCD, Na^+^, and Cl^-^ in coexisting dilute and dense phases as a function of total NaCl concentration, which we refer to as the total mean ion activity, and denote as *a*_NaCl_ (**Figure 2C, 2D** and **Figure S1B, S1C**). The data are shown as coexistence curves annotated by tie lines that connect points within coexisting phases. For a given tie line, the low and high concentration values along the horizontal axis correspond to activities of A1-LCD in the dilute and dense phases, respectively (**Figure 2C, 2D**). The measured activities of Na^+^ in dilute and dense phases determine the slope of the tie line, and each tie line corresponds to a specific value of *a*_NaCl_. The slopes of tie lines are negative when the coexistence curve is plotted on a plane defined by the input concentration of A1-LCD along the horizontal axis and input activity of Na^+^ along the vertical axis. Negative slopes point to preferential accumulation of Na^+^ in the dilute phase and preferential exclusion of Na^+^ from the dense phase. This sets up a positive Nernst potential V_N+_ and its magnitude decreases as the value of *a*_NaCl_ increases (**Figure 2E**). The implication is that application of a positive voltage to the dilute phase should drive additional sodium ions from the dilute phase into the dense phase, countering the preferential exclusion of sodium ions from the dense phase.

Preferential exclusion of Na^+^ (**Figure 2C**) from the dense phase is accompanied by preferential accumulation of Cl^-^ in the dense phase (**Figure 2D**). This also gives rise to a positive Nernst potential V_N-_ whose magnitude decreases with increasing *a*_NaCl_, crossing over to be negative at high values of *a*_NaCl_. The Nernst potential of chloride is positive because the charge of the ion is included in its calculation. Accordingly, exclusion of cations and accumulation of anions result in positive Nernst potentials for cations and anions (**Figure 1A**). Application of a positive voltage across the dilute phase should draw chloride ions out of the dense phase and into the dilute phase, thus countering the preferential accumulation of chloride in the dense phase. The Nernst potentials computed using direct measurements of interphase differences in Na^+^ and Cl^-^ activities may be interpreted as follows: For *a*_NaCl_ ≈ 200 mM, a potential of approximately 22 mV will balance the differences in Na^+^ activities across the coexisting phases. Likewise, for *a*_NaCl_ ≈ 200 mM, a positive potential of approximately 9 mV will balance the differences in Cl^-^ activities across the coexisting phases. This type of depolarization, achieved by applying an external potential drop, will affect the underlying phase behaviors that will likely be compensated either by changes to macromolecular concentrations in coexisting phases or dissolution of condensates, which will be reset once the balancing potential is removed.

The differential partitioning of cations and anions also sets up a Donnan potential, which we quantify using the Goldman-Hodgkin-Katz equation (**Figure 1A**). For A1-LCD condensates, the values of V_D_ decrease from ≈ +22 mV for *a*_NaCl_ ≈ 120 mM to ≈ +9 mV for *a*_NaCl_ ≈ 500 mM. This decrease arises from a widening gap between the Nernst potentials as *a*_NaCl_ increases and from asymmetries of *a*_NaCl_-dependent changes to the activities of Na^+^ and Cl^-^ in 1:1 electrolytes ^32^. Physically, a non-zero interphase Donnan potential implies that A1-LCD condensates are akin to mesoscale capacitors. The intrinsic capacitance is set by the macromolecule that drives phase separation, and the amount of charge that can be stored in the capacitor is determined by the magnitude of V_D_.

For A1-LCD, the driving force for phase separation is enhanced with increasing *a*_NaCl_. This salting-out behavior implies that the length of the tie line increases because the activity of A1-LCD in the coexisting dilute phase decreases with increasing *a*_NaCl_. In contrast, for a salting-in system, an increase in ion activity will cause a dissolution of the two-phase system. For the A1-LCD system, the interphase differences in cation-versus anion-specific Nernst potentials increase with increasing *a*_NaCl_, whereas the magnitude of the Donnan potential decreases with increasing *a*_NaCl_. The implication is that the increase in osmotic pressure that accompanies an increase in salt concentration, is balanced by lowering the macromolecular activity in the dilute phase and enhancing the flux (Nernst potentials) of ions across the two phases, thereby lowering the capacitance by discharging the condensate.

### Macromolecular charge affects the differential partitioning of Na^+^ and Cl^-^ ions

Donnan and Nernst potentials are determined by the overall electrochemical equilibrium of components between phases. Therefore, the macromolecular charge, the total mean ion activity, and the types of ions that make up the monovalent salt should alter Donnan or Nernst potentials. To test for this, we performed measurements by tuning the charge of the driver of phase separation, and measured dilute and dense phase activities of proteins as well as the activities of Na^+^ and Cl^-^ ions for condensates formed by A1-LCD and three different variants designated as -4D, +7K+12D, and +12D (**Figure 2B**) ^24^.

For the -4D variant, the net charge is positive (+12). In this variant, the four naturally occurring Asp residues were replaced with Gly / Ser (**Table S1**). For condensates formed by the -4D variant, we observe preferential accumulation of Na^+^ in the dilute phase and preferential exclusion of Na^+^ from the dense phase (**Figure 3A**). The Nernst potential V_N+_ is positive and it first increases and then plateaus as *a*_NaCl_ increases (**Figure 3C**). In contrast to Na^+^, Cl^-^ is preferentially incorporated within dense phases formed by -4D. This gives rise to positive slopes for tie lines (**Figure 3B**). The resultant Nernst potential remains positive at low values of *a*_NaCl_. The magnitude decreases with increasing *a*_NaCl_ and crosses over to being negative at high salt concentrations (**Figure 3C**). The Donnan potential is positive across the range of *a*_NaCl_ values that were studied (**Figure 3C**).

**Figure 3:**
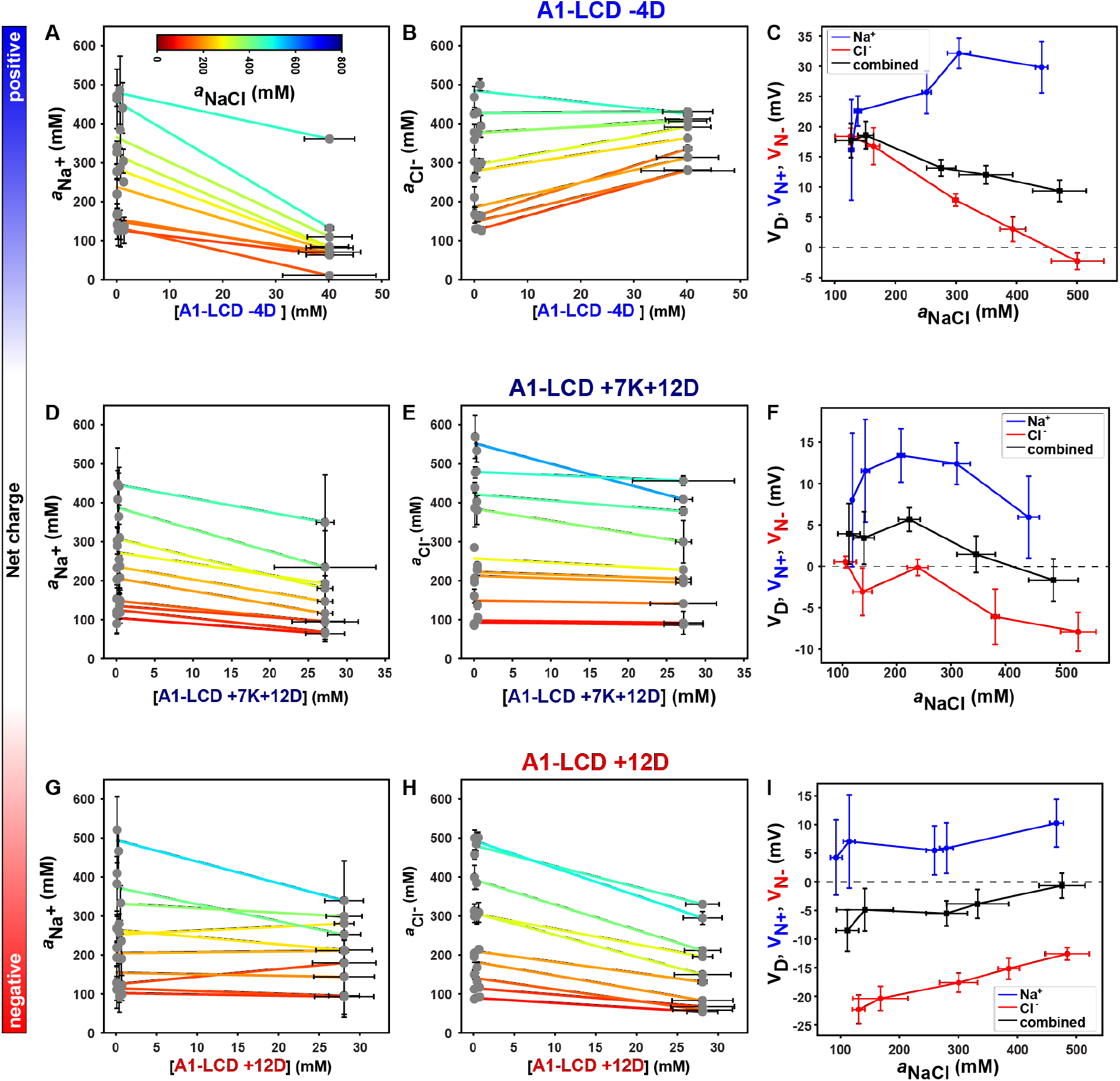
Interphase partitioning of cations and anions depends on protein charge. (**A, B, D, E, G, H**) Measured tie lines at different values of a_NaCl_ for the three A1-LCD variants including -4D (**A, B**), +7K+12D (**D, E**), and +12D (**G, H**). (**C, F, I**). For each variant, the interphase Nernst and Donnan potentials were computed using the values of the measured activities of Na^+^ and Cl^-^. **(C)** For the -4D variant, V_N+_, V_N-_, and V_D_ have positive values. The magnitude of V_N-_ decreases with increasing *a*_NaCl_, whereas the magnitude of V_N+_ first increases and then plateaus as *a*_NaCl_ increases. **(D)** For +7K+12D, the net charge is close to zero. Accordingly, V_N+_ and V_N-_ are positive and negative, respectively. The Donnan potential decreases and switches from being positive to being negative above *a*_NaCl_ ≈ 400 mM. **(E)** For +12D, which has a net negative charge, V_D_ is negative, and its magnitude decreases with *a*_NaCl_. The horizontal dashed lines that intersect the ordinate at values for potentials that are zero are drawn to guide the eye in panels **C, F**, and **I**. See **Figure S2** and **Figure S3** for additional data. As noted in **Figure 2**, tie lines that do not cross one another in 3-space can cross one another in 2-space. The tie lines plotted in 3-space make this point as shown in **Movies S2-S4**.

The +7K+12D variant has a net charge of +3. In this variant, nineteen Gly / Ser residues from A1-LCD WT were replaced with seven Lys and twelve Asp residues (**Table S1**). For condensates formed by the +7K+12D variant, the slopes of the tie lines for Na^+^ are negative for all values of *a*_NaCl_, demonstrating preferential exclusion of Na^+^ from the dense phase (**Figure 3D**). However, the slope of the tie line for Cl^-^ is close to zero for low to intermediate values of *a*_NaCl_, and switches to being negative for higher values of *a*_NaCl_ (**Figure 3E**). Accordingly, the Donnan potential switches from being positive to negative as *a*_NaCl_ increases (**Figure 3F**). Hence, there is a value of *a*_NaCl_ between 300 mM and 400 mM for which V_D_ ≈ 0. A zero value for V_D_ can arise if the activities of all ions are equal across the coexisting phases, i.e., 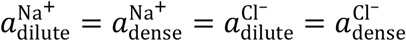. Alternatively, V_D_ can also be zero if the sum of 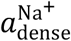 and 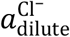 equals the sum of 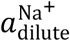 and 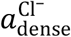, but 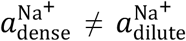 and 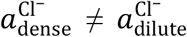.

Therefore, V_D_ can be zero if both V_N+_ and V_N-_ are zero or even if the Nernst potentials are non-zero. For the +7K+12D variant, the Donnan potential passes through zero even though V_N+_ remains positive for all values of *a*_NaCl_. In accord with the slopes of tie lines for Cl^-^ (**Figure 3E**), the Nernst potential V_N-_ switches to being negative for values of *a*_NaCl_ that are above 250 mM.

The net charge of the +12D variant is -4. Here, twelve Gly / Ser residues in A1-LCD WT were replaced by Asp residues. For low values of *a*_NaCl_ we observe minimal preferential exclusion of Na^+^ in the dense phase formed by the +12D variant. Beyond values of *a*_NaCl_ ≈ 300 mM, the tie line slopes become negative for Na^+^ (**Figure 3G**). We observe stronger exclusion of Cl^-^ from the dense phase, as evidenced by the negative slopes of tie lines for Cl^-^ (**Figure 3H**), which contrast with the mostly positive slopes of tie lines for A1-LCD WT (**Figure 2D**) and the -4D variants (**Figure 3B**). As a result, the Donnan potentials are negative for the +12D variant across the range of *a*_NaCl_ values that were titrated (**Figure 3I**). The Nernst potentials for Na^+^ are largely independent of *a*_NaCl_ and within error they are either weakly positive or close to zero. In contrast, the preferential exclusion of Cl^-^ from the dense phase weakens with increasing *a*_NaCl_ and this results in negative Nernst potentials V_N–_ for Cl^-^ whose magnitude decreases with increasing *a*_NaCl_ (**Figure 3I**).

Next, we asked if differential partitioning preserves electroneutrality of the entire system, and of each of the coexisting phases. We tested for electroneutrality by assuming and assigning charges to proteins based on their model compound pK_a_ values ^33^. Within the limits imposed by this assumption ^34^, we find that electroneutrality is preserved, as gleaned by summing the measured concentrations of protein charges and those of ions that partition into dense phases (**Figure S3**). Therefore, while we observe asymmetric ion partitioning across the phases, electroneutrality of the entire system and each of the coexisting phases is preserved.

### Interphase potentials depend on the type of ions that are in solution

Monovalent ions have different sizes, electronegativities, hydration free energies, numbers of water molecules in the first hydration shell ^35^, and lattice constants ^35-40^. A1-LCD forms condensates in a variety of monovalent alkali halide salts (**Figure S1A**). We quantified the interphase electric potentials based on different types of alkali halide ions with distinct chaotropic or kosmotropic properties ^38-40^. We hypothesized that different ions affect the solvation of IDPs differently. Accordingly, the measured asymmetry in ion partitioning should depend on ion types, thereby affecting interphase potentials of condensates.

We measured phase diagrams and the differential partitioning of monovalent cations and anions for A1-LCD condensates formed in NaBr, NaI, KCl, and LiCl (**Figure 4** and **Figure S4**). The slopes of tie lines clearly depend on the types of ions in solution. The impacts of ion sizes and charge densities of ions are inferred by comparing the signs and magnitudes of V_N+_, V_N-_, and V_D_ in NaCl (**Figure 2F**) versus NaBr and NaI (**Figure 4C, 4F**). Room temperature lattice energies are useful physical parameters that distinguish different alkali halide salts from one another ^36^. These energies measure the strengths of cohesive interactions of ions in monovalent salt crystals and are useful proxies for interpreting hydration free energies and for classification of ions and ion pairs into salting-in versus salting-out electrolytes ^36, 38^. Measurements of differential partitioning of cations and anions show that for sodium salts, the Nernst and Donnan potentials have similar signs and magnitudes for the low charge density iodide salt. Conversely, in NaBr and NaCl, V_N+_ and V_D_ are positive, whereas V_N-_ changes from positive to negative with increasing activity of the salt. These data show that the differential partitioning of solution ions and the interphase electric potentials they generate are tied to the charge densities and hence hydration properties of solution ions ^38-40^.

**Figure 4:**
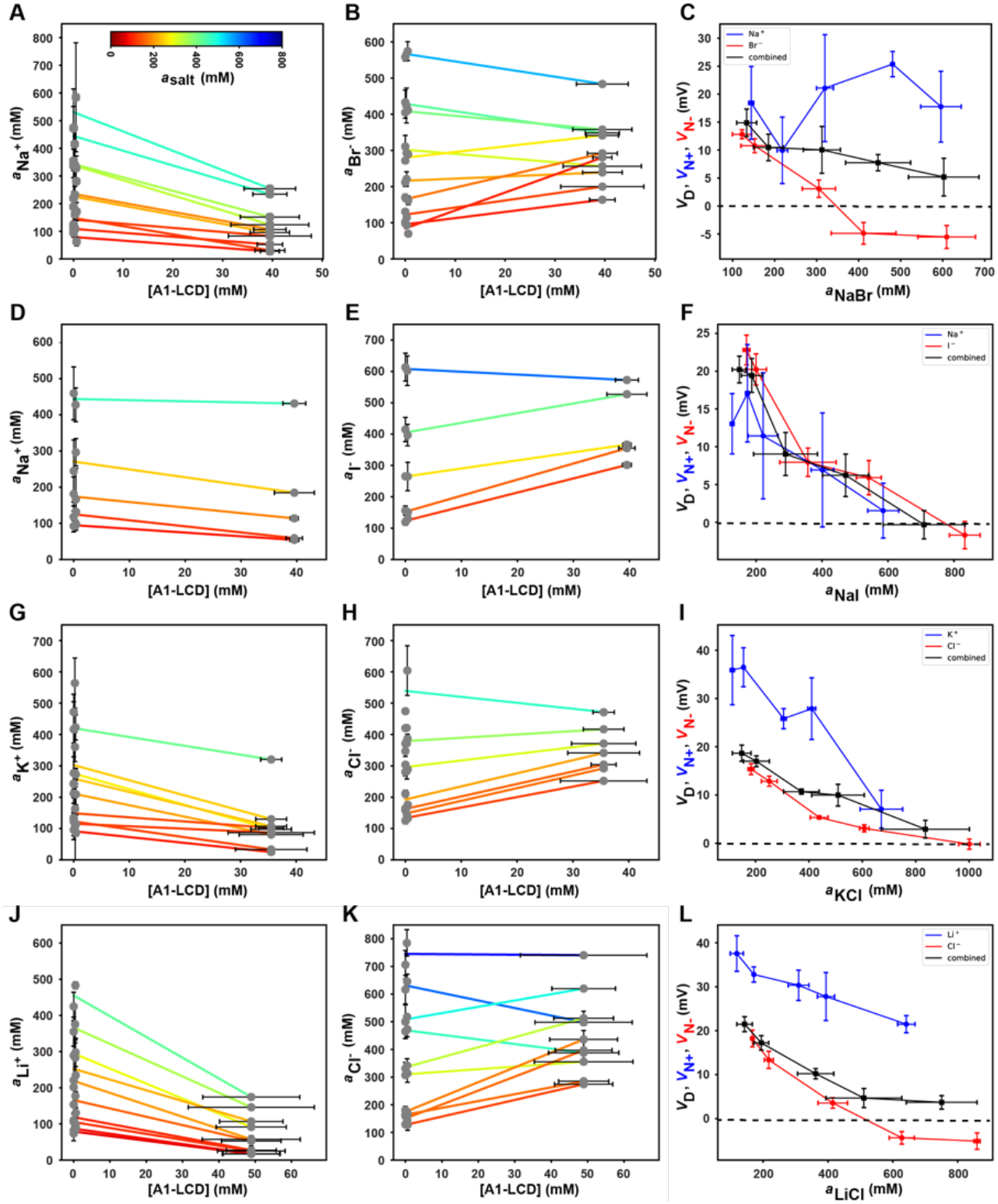
Interphase partitioning of ions varies with the types of ions in solution. Data are shown for tie lines measured for coexisting phases formed by A1-LCD WT in different amounts of NaBr (**A, B**), NaI (**D, E**), KCl (**G, H**), and LiCl (**J, K**). The Nernst and Donnan potentials computed using measured values for dense and dilute phase activities of cations and anions plotted against the total mean ion activities of the different salts are shown in (**C**) for NaBr, (**F**) for NaI, (**I**) for KCl, and (**L**) for LiCl. The data shown here are to be compared to the data shown in **Figure 2** for interphase measurements performed for A1-LCD WT in different concentrations of NaCl. In panels C, F, I, and L dashed horizontal lines are drawn to intersect the ordinate at potential values of zero. See additional data in **Figure S4**. As noted in **Figures 2** and **3**, tie lines that do not cross one another in 3-space can cross one another in 2-space. The tie lines plotted in 3-space make this point as shown in **Movies S5-S7**.

Ions can be classified as salting-in, or salting-out species based on their charges and free energies of hydration of ions (**Figure 5A**). We summarize the findings from characterizations of tie lines as well as Nernst and Donnan potentials (**Figure 4**) by plotting the variation of V_D_ against the room temperature lattice energies for each of the monovalent salts (**Figure 5B**). This comparison is performed by estimating the potentials for a total mean ion activity of 294 mM. We find a V-shaped dependence of the Donnan potentials on lattice energies, with NaBr located at the vertex of the V-shaped profile (**Figure 5B**). For A1-LCD condensates, salts that lead to increases or decreases in the magnitude of the lattice energies vis-à-vis NaBr are characterized by larger, more positive Donnan potentials. In contrast, for condensates formed by the +12D variant, the net charge is negative, and we observe an upside-down V-shaped profile for the variation of the Donnan potential with the lattice energies of the different salts. Again, NaBr is located at the vertex of the upside-down V-shaped profile. The Donnan potentials become increasingly negative as the lattice energies increase or decrease in magnitude, and they cross zero, becoming positive in the presence of NaBr. Such V-shaped behaviors have been reported in studies of correlations between standard heats of solution of crystalline alkali halide salts and differences between the absolute free energies of hydration of the ions ^39^. The Donnan potential is proportional to the amount of charge that can be stored in a condensate. The V-shaped profiles and salt type dependence of the Donnan potentials suggest that the amount of charge that can be stored is governed by differences in hydration free energies, lattice energies, and the ion associations in alkali halide solutions ^36, 39, 42^.

**Figure 5:**
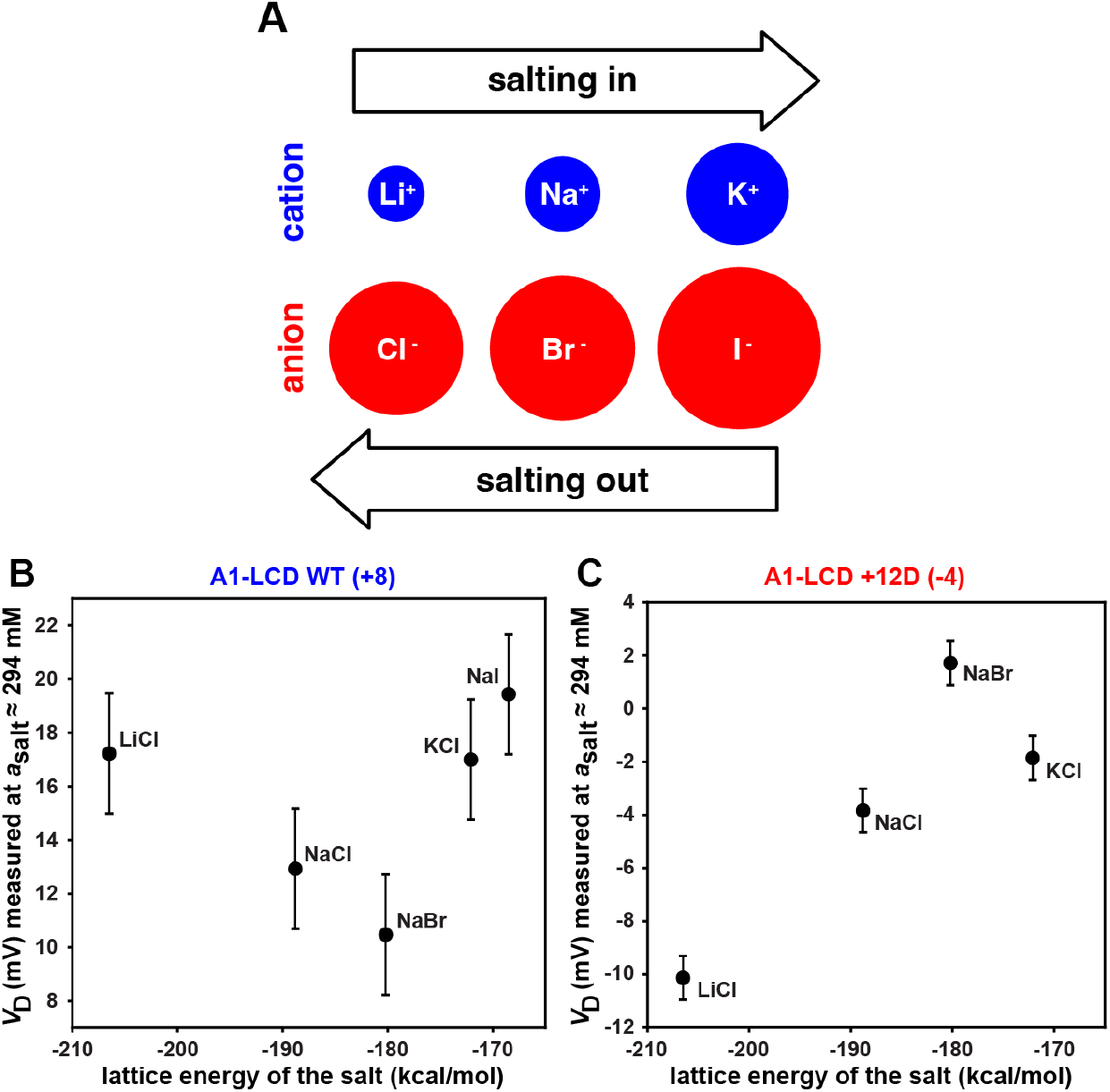
Donnan potentials are influenced by the types of ions that make up the monovalent salt. **(A)** Relative sizes of the salt ions used in this study are shown, as well as the relative salting-in vs. salting-out trends identified by Hofmeister ^41^. **(B)** The interphase Donnan potentials for A1-LCD WT vary with salt type for a fixed total salt concentration. The room temperature lattice energy of the salt is used as a representative quantitative feature with which to describe and compare salts ^36^. **(C)** The relationship between interphase Donnan potentials and salt lattice energy at a fixed total salt concentration is inverted when the net charge of the A1-LCD is inverted from +8 for A1-LCD WT to -4 for the +12D variant.

### Condensates formed by RNA molecules are also defined by interphase electric potentials

Most intracellular condensates are mixtures of proteins and RNA molecules ^6^. Recent studies have shown that RNAs can form condensates at high concentrations of monovalent salts ^25, 43^ or in the presence of divalent ions such as Mg^2+^ ^43, 44^. Furthermore, in RNP granules, RNA-RNA interactions appear to be key players in seeding ^45^ and stabilizing condensates ^45, 46^. RNA-driven condensation is likely to be a key driver of condensates in fungi and planta where the internal solution conditions are directly influenced by the environment ^47^. Therefore, as a model system, we investigated condensates formed by poly-rA in the presence of molar concentrations of different monovalent salts ^25^. These high concentrations of monovalent salts are needed for observing protein-free RNA condensation *in vitro*. In cells, the presence of multivalent ions including protein-based macroions enable RNA condensation at lower concentrations of monovalent salts. Polydisperse solutions of poly-rA form condensates in molar amounts of LiBr, NaBr, KBr, and NaCl (**Figure 6A**), and a small number of tie lines are accessible to direct measurements.

**Figure 6:**
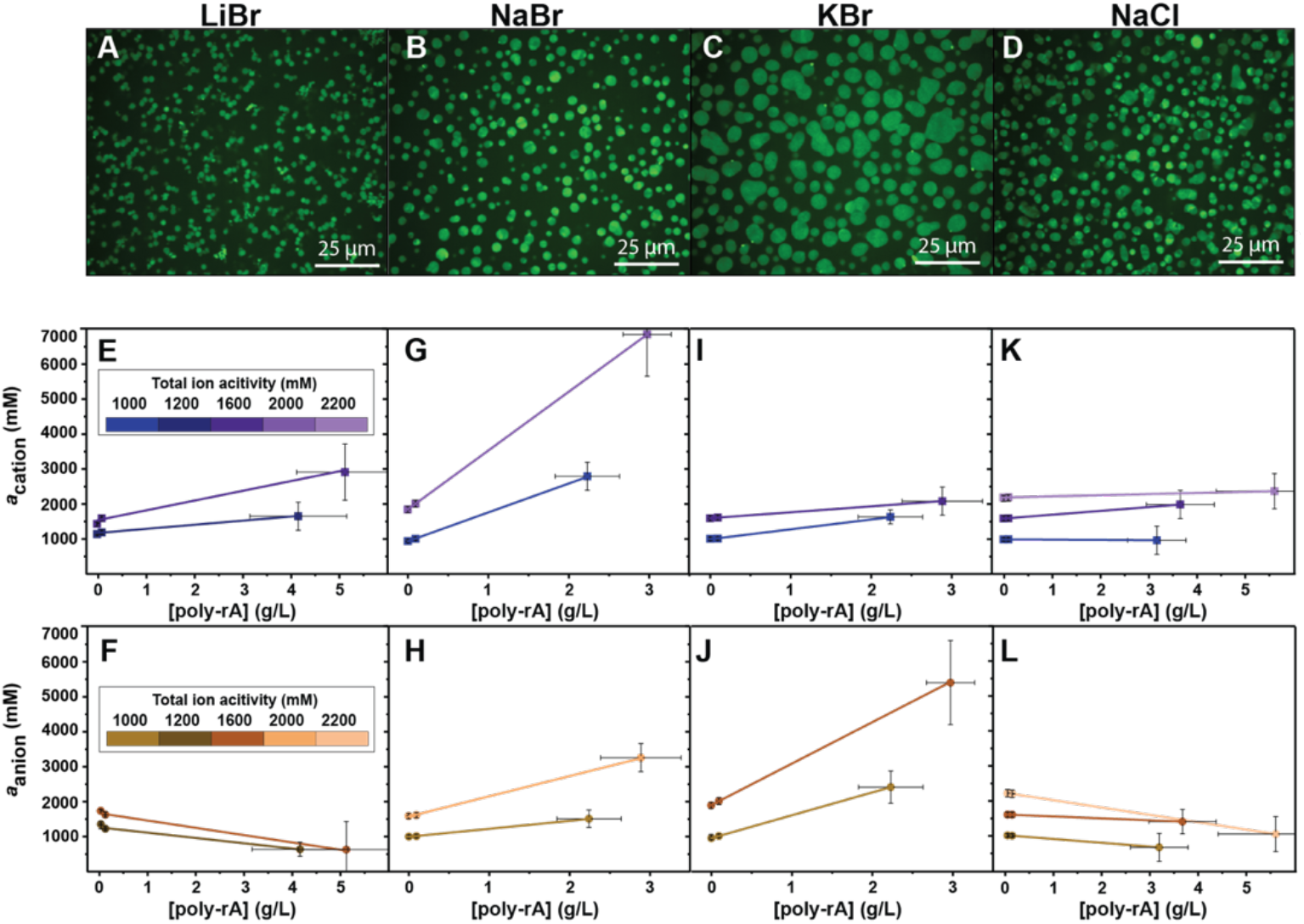
Differential partitioning of cations and anions in poly-rA condensates depends on ion types: Confocal images of poly-rA condensates in 1.4 M mM LiBr (**A**), NaBr (**B**), KBr (**C**) and NaCl (**D**). To enable imaging, 1% of the poly-rA molecules were labelled with Alexa Fluor647. We measured dilute and dense phase concentrations of poly-rA as well as cations and anions at different total mean ion activities – see color bars in panels (**E, F**). Results from the measurements are shown as coexistence curves mapped onto planes showing the measured poly-rA concentrations along the horizontal axes in panels (**E**-**L**), the measured activities of cations and anions along vertical axes for LiBr (**E, F**), NaBr (**G, H**), KBr (**I, J**), and NaCl (**K**), (**L**). Columns going from left to right show data for poly-rA condensates in LiBr, NaBr, KBr, and NaCl.

In bromide salts of Li^+^, Na^+^, and K^+^, we observe preferential accumulation of cations in the dense phase over the dilute phase (**Figure 6B-6I**). This makes intuitive sense since the accumulation of cations is essential for neutralizing the negative charge on the polyphosphate backbone of poly-rA. Interestingly, Br^-^ ions are not always preferentially excluded from the dense phase. While the slopes of tie lines for Br^-^ are negative in LiBr, they are positive in NaBr and KBr (**Figure 6B-6I**). This points to the accumulation of co-ions (Br^-^) alongside accumulation of counterions (Na^+^ or K^+^) in dense phases formed by poly-rA. These results are reminiscent of ion-type and ion activity dependent accumulation versus exclusion of counter- and co-ions ^48^ that have been observed for RNA molecules ^49^.

In NaCl, as opposed to NaBr, we observe positive or near zero slopes of tie lines for Na^+^ (**Figure 6K**) and negative slopes of tie lines for Cl^-^ (**Figure 6L**). The measurements of interphase ion activities allow us to quantify interphase Nernst and Donnan potentials (**Figure 7**). In LiBr, the interphase Donnan potentials are negative or close to zero, as is the Nernst potential for Br^-^ (V_N-_), while the Nernst potential for Li^+^ (V_N+_) is positive. In NaBr and KBr, the Donnan potentials as well as the Nernst potentials are positive, and they have a larger magnitude for NaBr than for KBr. In NaCl, the Donnan potentials are close to zero, and the Nernst potentials are either positive (V_N+_) or negative (V_N-_). This shows that a near zero membrane-like potential can still involve an asymmetry in cation / anion accumulation or exclusion (**Figure 7**).

**Figure 7:**
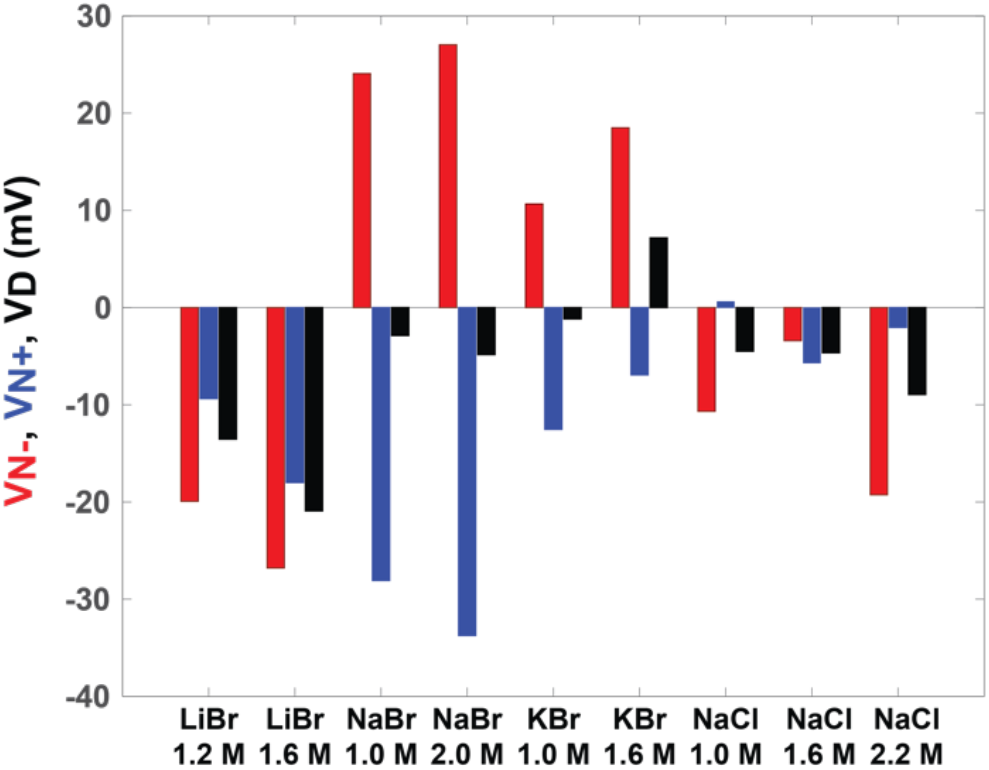
Interphase potentials of poly-rA condensates. Nernst and Donnan potentials computed using direct measurements of cation and anion activities in dilute and dense phases of poly-rA condensates.

## SUMMARY

### Implications of condensates being defined by interphase electric potentials

Our data suggest that condensates, which are defined by coexisting dense and dilute phases, are akin to mesoscale capacitors. The intrinsic capacitance will be set by the net charge of macromolecule(s) that accumulate asymmetrically across dense and dilute phases. The Donnan potential determines the amount of charge that can be stored in condensates. Note that capacitors can pass alternating currents but not direct currents. This is because they are repeatedly charged and discharged as the direction of current flow changes. The implication is that condensates can enable the starting of molecular and mesoscale engines such as induction motors that cannot self-start, and therefore do work. An example of a condensate that has been shown to do work to invaginate membranes comes from yeasts where condensates formed by the PLCD of Sla1 enables actin-independent vesicle formation ^50^. Further, such capacitor effects suggest that perturbations of interphase electrochemical equilibria should result in a reorganization of ion flux. This can serve as modulators of ion motive forces ^10^.

The interphase potentials help rationalize recent findings regarding the electrochemical impacts of condensation on cellular behaviors. Dai et al., showed that condensate formation has a direct effect on intracellular electrochemical properties ^51^. These include cytoplasmic pH and hyperpolarization of the membrane potential. The effects, measured in bacteria, were conjectured to be due to condensates being defined by the presence of interphase electric potentials. Here, we show this to be true by quantifying interphase potentials using direct measurements of ion activities in coexisting phases. Our findings suggest that modulation of the interphase potentials, which we show to be influenced by macromolecular charge, ion type, and salt concentration, might explain the specificity of uptake of small molecules by cells. This, as shown by Dai et al., has a direct impact on bacterial survival under antibiotic stress ^51^. Dai et al., also demonstrated that shifts of the intracellular electrochemical equilibria, driven by condensate formation, have a profound influence on gene expression profiles ^51^. All these effects can be rationalized by the interphase electric potentials that we measure and report in the current work.

Interphase potentials will help set up an electric double layer at the interface of condensates. These interfacial potentials have been measured as zeta potentials in condensates formed by FUS and poly-rU ^52^, as redox potentials in condensates formed by resilin-like polypeptides ^53^, and as strong interfacial electrical responses of condensates formed by synapsins ^54^. The Gibbs dividing surface or plane ^55^ is situated at the interface, and it delineates the boundary between the coexisting phases. An interphase ion gradient must be balanced at the interface, and hence, in addition to the interphase potential gradient, there will also be a potential gradient between the coexisting phases and the interface. This electric potential gradient can set up a potential barrier or potential well that influences passive molecular transport. Our data show that the interphase potential can be positive or negative, and this suggests that the directionality of the interfacial electric potential can also be varied by modulating the differential partitioning of ions across coexisting phases. Directional potentials prescribe the direction of passive molecular transport of charged species, and this can serve as the physical basis for understanding the intrinsic roles of condensates as regulators of cell signaling and controllers of macromolecular fluxes ^51, 56^.

Under osmotic stress, such as hypertonicity, cells lose water, and their survival depends on being able to recover cellular volume to survive ^57^. A recent study showed that heat shock can influence the activity of water in cells, and water potentials are regulated by condensation of IDRs. Our studies, which focus on direct measurements of interphase potentials, help provide a physical basis why and how condensates can buffer water or ion potentials ^57^. For example, in renal cells, recovery of cellular volume is enabled by WNK1 kinases that form condensates ^58^. This initiates phosphorylation-dependent signaling to enable a net influx of ions to help restore cell volume. WNK1 condensation was found to be driven by its C-terminal IDR, and this domain was postulated to function as a crowding sensor ^58^. Based on our work, we propose that the WNK1 condensates play two roles. The osmotic equilibrium will be influenced by changes to water or ion abundance, and this will alter interphase electric potentials in response to osmotic stresses. The second role pertains to the kinase activity of WNK1. We propose that the salt-concentration-dependent interphase potentials of WNK1 condensates enable their functions as electrochemical sensors, and regulators of osmotic stress. Our work, taken together with that of Boyd-Shiwarski et al.,^58^ suggests that the tunability of interphase electric potentials might serve as a target for diagnostics and therapeutics aimed at controlling cell volume changes in renal cells. Other applications in cell physiology, pathology, and synthetic biology are likely to be forthcoming as we gain a deeper appreciation of the broader implications of interphase electric potentials that define biomolecular condensates.

Our work highlights the importance of direct measurements of ions, solutes, and metabolites in coexisting phases. Future investigations will focus on adapting our methods to enable measurements of interphase potentials and their functional consequences in cellular milieus. This will require using electrochemical probes for in situ measurements in live cells. The latter will require adaptations such as direct measurements based on plasmonic nano-sensors ^59^ or fluorescent-based environmentally sensitive reporters ^10, 11, 60^.

## METHODS

### Materials

Poly-rA (MW 700-3500 kDa, P10108626001), 70% nitric acid, Sodium phosphate dibasic and monobasic, citric acid, HEPES, Tris, CAPS, LiBr, NaBr, KBr, NaCl and KCl were obtained from Sigma Aldrich. RNAase free water was purchased from Thermo Fisher.

### Proteins used in this study

We used the wildtype prion-like disordered low-complexity domain (residues 186-320) of the human RNA-binding protein hnRNPA1 (UniProt: P09651; Isoform A1-A). Additionally, we included three variants, in which we replaced glycine and serine residues with 12 aspartate residues (+12D), or 7 lysine and 12 aspartate residues (+7K+12D), or we removed the 4 aspartate residues (−4D) and replaced them with glycine and serine residues that were present in the WT A1-LCD sequence. We used the same purification strategy as previously described by Bremer et al. ^24^. The purified protein samples were stored in 6 M GdmHCl (pH 5.5), 20 mM MES at 4°C until usage. The identity of each protein was verified via intact mass spectrometry. Amino acid sequence details for each of the constructs are shown in Supplementary Table 1 below. All protein constructs contain the two additional residues “GS” at the N-terminus (italic), which remain after cleavage with TEV protease is performed.

### A1-LCD sample preparation for ICP-MS and ISE

Protein solutions in denaturant were exchanged into native buffer as previously described ^24^. Phase separation was induced by adding the specified concentration of salt to the protein sample in 20 mM HEPES (pH 7.0). This solution will be referred to as the primary stock. 2 μL of the total primary stock was pipetted into 500 μL neat (70%) nitric acid (Trace Metal Grade, Fisher Scientific) in a 10 × 75 mm borosilicate glass tube (Fisher Scientific) for subsequent boiling (see below). This is the “total” ICP-MS sample. In parallel, 2 μL of the primary stock was diluted 1:1000 in ddH_2_O for subsequent anion concentration determination using an ion selective electrode (ISE) for the anion of interest (“total” ISE sample). After the 2 μL total samples were taken from the primary stock, the primary stocks were incubated for at least 20 min at 4°C to allow for phase separation. The samples were then centrifuged for 10 min at 12,000 rpm to separate dilute and dense phases. 2 μL each of dilute phase and dense phase were then pipetted into separate 10 × 75 mm borosilicate glass tubes with 500 μL neat (70%) nitric acid as described above (“dilute phase” and “dense phase” ICP-MS samples). Another 2 μL each of the dense and dilute phases from the primary stock were diluted 1:1000 in ddH_2_O for subsequent ISE measurement (“dilute phase” and “dense phase” ISE samples). A glass marble was placed on each of the glass tubes to prevent evaporation during heating, and the samples were boiled for 1 h at 110°C on a heating block. After cooling, samples were further diluted in two steps prior to ICP-MS measurements: First, each 500 μL boiled sample was diluted into 34.5 mL ddH_2_O. Second, 500 μL was pipetted from the 35 mL step 1 dilution and diluted into 2000 μL 1% nitric acid (previously prepared and filtered with a 2 μm PES filter).

Dense phase samples were pipetted with a positive displacement pipette. In parallel, we determined the protein concentrations in coexisting dilute and dense phases using classic UV absorbance with a spectrophotometer, or analytical HPLC when the absorbance at 280 nm was below 0.1. Dense phase protein concentrations are plotted as the average across all the salt concentrations in each dataset. The error bars in the concentration dimension for dense phase samples represent the absolute deviation from this average. This was justified given that the dense phase concentration is not expected to change significantly with salt concentration and the variation that we did observe in measurements of dense phase concentration across salt concentration appeared to be random. All solutions were prepared with ultrapure water (18.2 MΩ cm of resistivity) from a Milli-Q Integral 3 purification system (Millipore, Bedford, MA, USA).

### Poly-rA sample preparation for ICP-OES and IC

Poly-rA samples were prepared by mixing nuclease free water, poly-rA, optionally buffer and lastly salt solution. The stock solutions comprised poly-rA (30000 – 100000 ng/μL), LiBr (10 M), NaBr (7.5 M), KBr (5.6 M), NaCl (6 M), 100 mM phosphate (pH 2.5), 100 mM citrate (pH 3.3, 4.2, 5.2, or 5.7, respectively) and 100 mM HEPES (pH 7.1 or 8.1, respectively) in nuclease-free water. After condensate formation, dense and dilute phases were separated by spinning down at 14,000 x g for 10 minutes. The dilute and dense phase RNA concentrations were measured by Nanodrop. Then, the dilute or dense phases were diluted in 15 mL water to a maximum of 50 ppm. 5 mL of sample was used for the IC measurements. To 8 mL of sample, 235 μL 70% nitric acid was added for ICP-OES measurements. Additionally, blanks without salt and samples with known salt concentrations for calibration were prepared in a similar manner. Calibration curves were constructed with five samples or more. A blank and one calibration sample was run at least every 15 samples.

### ICP-MS measurements

ICP-MS measurements were carried out on a NexION 2000 inductively coupled plasma mass spectrometer (PerkinElmer, Waltham, MA, USA) equipped with a dynamic reaction cell (DRC) and autosampler (PerkinElmer). The instrument was run through a SmartTune procedure before each batch of measurements to ensure optimum functionality. A six-point standard curve from 200 ppb to 1 ppb (200, 100, 50, 10, 5, 1) was prepared from elemental standards (Perkin Elmer), plus a 0-ppb blank. To monitor drift, the 50-ppb standard was remeasured after every 6-10 samples. The raw data for total sample, dilute and dense phases were measured in counts per second (cps). These quantities were first converted to units of ppb using the standard curve, then converted from ppb to molar units, then back-calculated to the undiluted concentration by accounting for all dilutions made throughout the sample preparation process. Finally, the buffer blank (converted to molar units and back-calculated to account for dilutions in the same manner) was subtracted from each data point. Data for at least two biological replicates, prepared from different protein preparations and measured on separate days, were collected for each combination of construct and salt type (“batch”), and each batch consisted of samples spanning a range of salt concentrations, and with each salt concentration sample prepared in triplicate (technical replicates). To further assess and account for instrument variability, a 20-ppb scandium internal standard was introduced into each sample during auto-sampling for all measurements.

### Ion selective electrode (ISE) measurements

To determine anion and cation concentration, 2 μL of the total, dilute phase and dense phase were diluted 1:1000 with ddH_2_O in 15 mL falcon tubes. Bromide (ThermoScientific), chloride (Vernier), and iodide (Cole-Parmer) anion ISEs were used. Additionally, cation ISEs were used for sodium (Cole-Parmer) and potassium (Vernier). The LabQuestMini (Vernier) sensor interface was used for data collection for the Vernier ISEs. For all other ISEs, an AB15 Plus meter (Accumet) was used.

A five-point standard curve was measured with freshly prepared standard solutions before every batch of sample measurements to ensure precise measurements and accurate calibration of the ISE. The standard curve was fit to either an exponential (*y* = *be*^*Ax*^) or a quadratic equation to convert the values measured for unknowns to units of molar concentration. As with the ICP-MS measurements, a buffer blank was subtracted from each data point. 1:50 dilution of ionic strength adjuster (Thermo Fisher) was added to each sample for the bromide and sodium ISE. For each sample we measured at least two biological replicates, and for each biological replicate we measured 1-2 technical replicates of the dense phase sample, and 3 technical replicates of the total and dilute phase samples. Protein stock solution in 20 mM HEPES (pH 7.0) were measured as controls.

For each salt type, activity coefficients of electrolytes vs. electrolyte concentration data, as reported in The Handbook of Chemistry and Physics ^32^ were fit to fourth-order polynomials and the resulting fits were used to convert measured ion concentrations to activities.

### ICP-OES measurements

The concentrations of lithium, potassium, and sodium ions in poly-rA samples were determined using a Thermo Fisher Scientific iCap 7400 Duo inductively coupled plasma optical emission spectrometer (ICP-OES). 10 ppm scandium was used as the internal reference and samples were run three times. The absorption at 610.362 nm (lithium), 589.592 nm (sodium) and 766.490 (potassium) was used to quantify concentrations.

### IC measurements

A Thermo Fisher scientific ion chromatography machine with suppressor (ADRS 600 4 mm), dionex column (Ionpac AS11 4 × 250 mm) and guard column (RFIC 4×50 mm) was used to measure the concentrations of chloride and bromide in poly-rA samples. Samples were loaded using a 25 μL loop and eluted over the column isocratic at 1 mL/mL at 12 mM KOH with the eluent generated from a Dionex eluent generator cartridge (EGC 500 KOH) for 5 minutes at 30 °C. At these settings, chloride and bromide eluted at approximately 2.1 and 2.8 minutes. The area under the curve of the peaks in the conductivity chromatogram was used to quantify concentrations.

Intensities (ICP-OES) and area under the curve (IC) were converted to concentrations of the undiluted samples using the calibration curves. Salt concentrations were then converted to estimated activities using reported mean activity coefficients of electrolytes at 25 °C as a function of concentration ^32^.

### Determination of protein concentrations in coexisting dense and dilute phases

Concentrations in coexisting dense and dilute phases were determined via UV-Vis spectroscopy except for A1-LCD WT. For this system, we used analytical HPLC to determine the protein concentration in the coexisting dilute phase because the absorbance at 280 nm was below 0.1 in these samples. 100-400 μL of the dilute phase were injected onto a ReproSil Gold 200 C18 column (Dr. Maisch) on a Waters HPLC system with a dual-channel UV/Visible Detector (Waters 2489). The column was equilibrated in 95% buffer A (ddH_2_O + 0.1% TFA; Sigma-Aldrich) and 5% buffer B (acetonitrile; Alfa Aesar). The protein was eluted from the column by running a gradient from 20-80% of buffer B over 5 column volumes.

In parallel, we measured a standard curve by injecting 5 different volumes of A1-LCD WT with known concentrations. The standard curve was used to determine the concentrations of the injected samples with known volumes ^26^. Dense phase protein concentrations on tie line plots are reported as the mean of all (ten) dense phase samples, and the horizontal dense phase error bars represent the absolute deviation from the mean.

### Quantification of Nernst and Donnan potentials

Interphase Nernst potentials were calculated using the Nernst equation, which is typically used to quantify the potential of a single ionic species across a membrane:

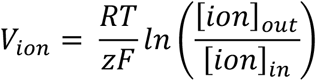

Here, *V*_*ion*_ is the Nernst potential for a given cation or anion, *R* = 8.314 J/K-mol is the ideal gas constant, *T* is the temperature in Kelvin, *z* is the charge number or ion valency (all salts used in this work were monovalent, so *z* = 1 for cations and *z* = -1 for anions), and *F* = 9.645 × 10^4^ C / mol is the Faraday constant. Applied to membraneless condensates, we consider [*ion*]_*out*_ to refer to the measured concentration (activities) of ions in the dilute phase, and [*ion*]_*in*_ to refer to the concentration of ions in the dense phase. Using these designations, we quantify the Nernst potentials as follows:

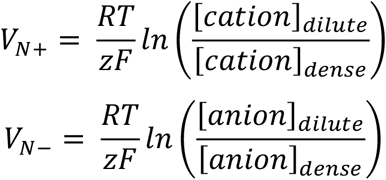

The values used for [*cation*]_*dense*_ and [*cation*]_*dilute*_ in the equation above were dense and dilute phase activities derived from concentration measurements made by ICP-MS, and the values used for [*anion*]_*dense*_ and [*anion*]_*dilute*_ refer to dense and dilute phase activities derived from concentration measurements made using ISE.

Interphase Donnan potentials were calculated using the Goldman-Hodgkin-Katz (GHK) equation, which is derived from the Nernst equation and accounts for all ionic species (anions and cations) present on either side of the phase boundary to determine the equilibrium interphase electrical potential across an interface. The equation also considers the relative permeability (*P*) of each ion:

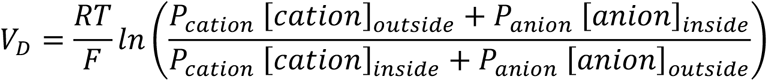

Again, we adapt this for membraneless condensates by designating *outside* as dilute phase and *inside* as dense phase. Because we are dealing with a phase boundary rather than a physical barrier, as would be the case for a lipid membrane, we set the permeability of all ions to be one, and this simplifies the GHK equation to the form shown in **Figure 1A** and reproduced below:

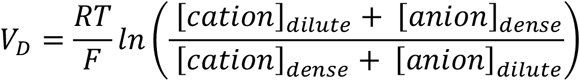

For calculations of dense phase net charge (condensate electroneutrality, **Figures S3** and **S4**) errors were calculated as the root squared sum (RSS) of the standard deviation of each of the components contributing to net charge:

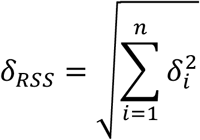

For the calculated Donnan potentials (**Figures 2-5**) the RSS error propagation takes the following form:

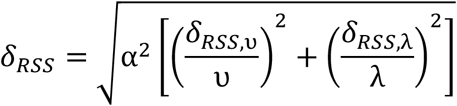

Here,

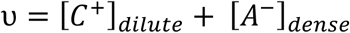

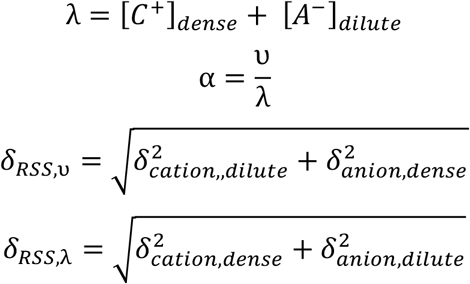

[*C*^*+*^]_*dilute/dense*_ and [*A*^*-*^]_*dilute/dense*_ are the measured concentrations of cations and anions in dense and dilute phases, and 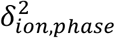 are the squared standard deviations of the corresponding concentration measurements for each ion in each phase.

### Software

Figures 1, 2, 3, 4 were made using python 2.7, Figures 5 and 6 were made using OriginPro 2017. Adobe Illustrator 2020 was used to organize panels of all figures.

## ASSOCIATED CONTENT

The Supporting Information includes **Figures S1-S4** and **Table S1**.

## AUTHOR INFOMRATION

### Author Contributions

Conceptualization: AEP and RVP; Methodology: AEP, NAE, AB, AP, TM, and RVP; Investigation: AEP, NAE, AB, AP, and RVP; Analysis: AEP, AB, NAE, RVP; Funding acquisition: TPJK, TM, and RVP; Project administration: TM, TPJK, and RVP; Supervision: YD, TM, and RVP; Writing: RVP; Review & editing: all authors

### Notes

RVP is a member of the scientific advisory board and shareholder of Dewpoint Therapeutics Inc. The other TPJK is a co-founder and chief technology officer of Transition Bio Inc. AEP is currently employed as a senior scientist at Auragent Bioscience in St. Louis, MO, USA. The other authors do not have any financial interests to declare.

## ACKNOWLEDGMENTS

We thank Elaine Flynn and Jeffrey Catalano for help with the use of the ICP-OES and IC, and Matthew King and Min Kyung Shinn for helpful discussions. This work was funded by the US Air Force Office of Scientific Research grant (FA9550-20-1-0241 to RVP), the St. Jude Research Collaborative on the Biology and Biophysics of RNP Granules (to TM and RVP), the US National Science Foundation (MCB-2227268 to RVP), the US National Institutes of Health (R01NS121114 to TM and RVP), the Royall Scholarship (to NAE), the Center for Biomolecular Condensates in the James McKelvey School of Engineering at Washington University in St. Louis (to AEP and NAE), the American Lebanese Syrian Associated Charities (to TM), and the European Research Council (ERC grants PhysProt agreement no. 33796 to TPJK).

