## Supplemental Materials for "Biomolecular condensates are characterized by interphase electric potentials"

[§]Corresponding Authors

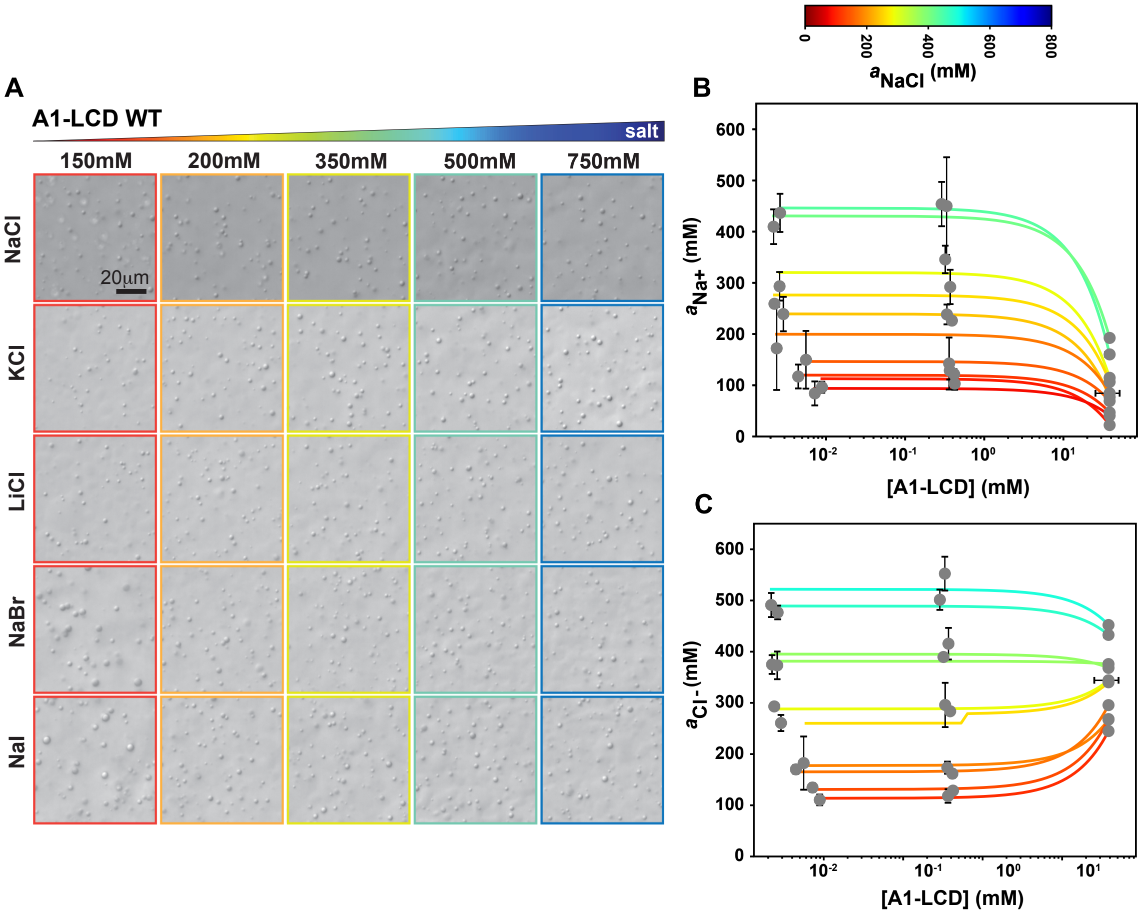


**Figure S1: A1-LCD forms condensates in different salts. (A)** Multi-panel plot of images from differential interference contrast (DIC) microscopy showing that A1-LCD forms condensates across a range of salt concentrations for five different alkali-halide salts. **(B)** and **(C)** Phase boundaries on the planes of protein concentration and measured ion activities in dense and dilute phases are shown in semi-log scale. Tie lines that have positive or negative slopes on linear scales will be curved in log scale. The semi-log scale helps establish the range of protein concentrations spanned by the system preparation, the points in the middle of the tie lines, and the values we measure for coexisting dense and dilute phases.

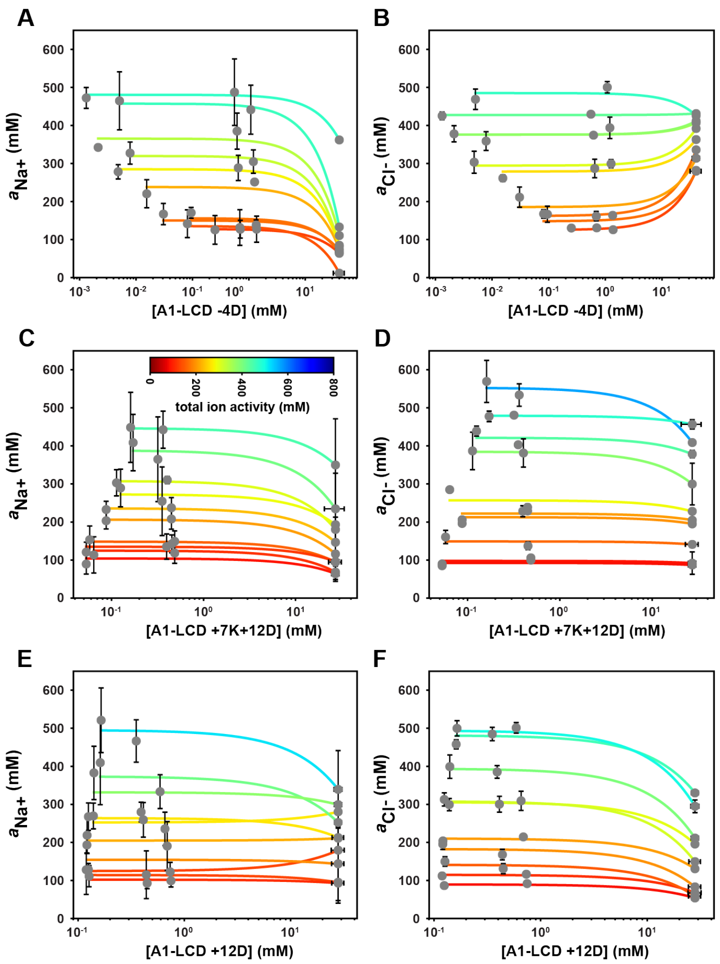


**Figure S2: Coexistence curves shown on semi-log scales, with ion activities in linear scale and protein concentrations on log scales.** Also see **Figure S1**. These are the same data as shown in Figure 3.

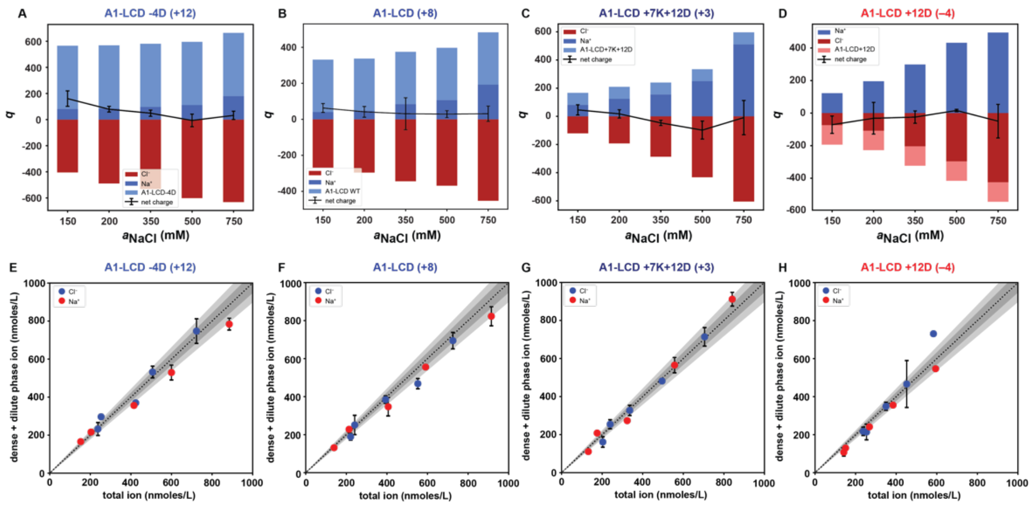


**Figure S3: Tests for electroneutrality and quantification of mass balance of the solution ions in NaCl.** (**A-D**) For each A1-LCD variant, we tested if the net charge within the condensate remains near zero at all total mean ionic activities of NaCl. For all ionizable species, we compute the charge by assuming the $pK_a$ values of model compounds [1]. We also assume the dense phase pH to be equivalent to that of the dilute phase. The sum of positive and negative charges is plotted as a line, with error bars representing the root sum of squared standard deviations for each of the components ($Na^+$, $Cl^-$, and protein concentrations). Deviations from a net charge (*q*) of zero are likely the result of charge regulation reflecting proton uptake or release. (**E-H**) The panels in the bottom row quantify the sum of measured ion activities in the dense and dilute phases along the ordinate and plot this against the concentrations of ions in the system at the start of the experiment. We observe a clear 1:1 correspondence, thereby showing that all solution ions have been accounted for in the measurements.

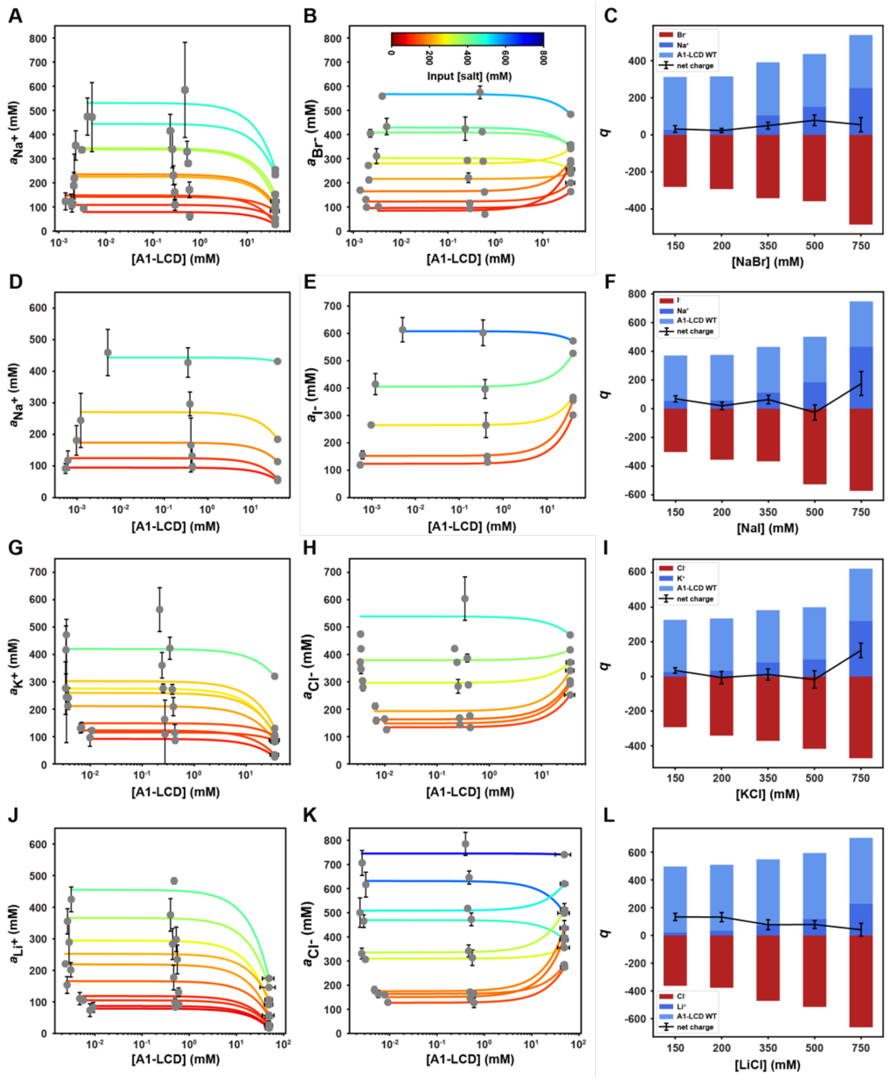


**Figure S4: Semi-log plots of tie lines and electroneutrality plots for condensates formed by WT A1-LCD in various salts. (A, B, D, E, G, H, J, K)** show tie lines cations and anions on the semi-log scale show the curvature of the dilute phase arm of the binodal and enable resolution of input and dilute phase data points. These are the same data as shown in Figure 4A-K. **(C, F, I, L)** For each salt type, we used the measurements and estimates of protein charge to compute net charge within the condensate. These estimates remain near zero at all salt concentrations when charge from salt ions and macromolecule are considered together, thus showing that electroneutrality is essentially always preserved. The sum of positive and negative charges is plotted as a line, with error bars representing the root sum of squared standard deviations for each of the components (cation, anion, and protein concentrations). At some salt concentrations there appears to be a slight net charge in the dense phase, which might be accounted for by charge regulation in the protein (e.g., neutralization of a positively charged residue via a $pK_a$ shift). Overall, the net charge appears to be small and somewhat random with respect to total salt concentration, so is likely to be due to the error of the measurement and lack of a full accounting for charge regulation effects and the effects of buffer ions.

**Table S1: Amino acid sequences of A1-LCD and designed variants**

| Construct | Amino acid sequence |
|---|---|
| A1-LCD | *GS*MASASSSQ RGRSGSGNFG GGRGGGFGGN DNFGRGGNFS GRGGFGGSRG GGGYGGSGDG YNGFGNDGSN FGGGGSYNDF GNYNNQSSNF GPMKGGNFGG RSSGPYGGGG QYFAKPRNQG GYGGSSSSSS YGSGRRF |
| A1-LCD +12D | *GS*MASADSSQ RDRDDSGNFG DGRGGGFGGN DNFGRGGNFS DRGGFGGSRG DGGYGGDGDG YNGFGNDGSN FGGGGSYNDF GNYNNQSSNF DPMKGGNFGD RSSGPYDGGG QYFAKPRNQG GYGGSSSSSS YGSDRRF |
| A1-LCD +12D+7K | *GS*MASADSSQ RDRDDKGNFG DGRGGGFGGN DNFGRGGNFS DRGGFGGSRG DGKYGGDGDK YNGFGNDGKN FGGGGSYNDF GNYNNQSSNF DPMKGGNFKD RSSGPYDKGG QYFAKPRNQG GYGGSSSSKS YGSDRRF |
| A1-LCD -4D | *GS*MASASSSQ RGRSGSGNFG GGRGGGFGGN GNFGRGGNFS GRGGFGGSRG GGGYGGSGGG YNGFGNSGSN FGGGGSYNGF GNYNNQSSNF GPMKGGNFGG RSSGPYGGGG QYFAKPRNQG GYGGSSSSSS YGSGRRF |
